# How fast can PPRV spread? Analyzing the results of an experimental PPRV infection of goats equipped with Ultra-WideBand sensors

**DOI:** 10.1101/2025.07.21.665851

**Authors:** Manal Nhili, Jean-Baptiste Menassol, Gaye Laye Diop, Mariame Diop, Aminata Ndoye, Mbengué Ndiaye, Aminata Ba, Michel Dione, Assane Gueye Fall, Noha El Khattabi, Modou Moustapha Lo, Mounia Abik, Andrea Apolloni, Adama Diallo

## Abstract

Peste des Petits Ruminants (PPR) is a highly contagious disease affecting goats and sheep. The speed and extent of its spread depend on contact patterns and the virus’s intrinsic characteristics. To estimate propagation speed and the role of inter-individual patterns, we analyzed the results of a series of 12 experimental infections conducted in five different sessions, involving six or seven goats in a secured stable. In each experiment, one animal was inoculated with the PPR strain and placed in contact with the other naïve animals. All the animals were equipped with Ultra-WideBand sensors to collect inter-individual distance data. The duration of this phase varied from 1 to 48 hours across the experiments. Afterwards, the animals were isolated and monitored for three to five weeks. Temperature, symptoms, nasal and ocular discharges, and blood samples were routinely collected to detect the presence of the virus. Using Bayesian statistical analysis, data on inter-individual distances, and health status were analyzed to estimate R_0_, the incubation period, and PPRV transmissibility. The latter was used to estimate the exposure period, i.e., the minimum amount of time for a naïve animal to become infected. Of the 70 naïve animals exposed to the virus, 18 were infected. R_0_ was estimated to be around 4.3 or 8.6, depending on the infectious period value. The incubation period was estimated to be around 16.3 days (95% CI: 12.6–23.3 days). The exposure time varies greatly depending on the density, ranging from nearly two days at low density to around four hours at high density. Large gatherings of animals, such as at livestock markets, could greatly facilitate the spread of PPRV. Furthermore, the long incubation period coupled with livestock mobility could favor the virus geographical dissemination on a large scale.

**Author summary:** PPR is an infectious disease that is transmitted directly and affects goats and sheep. Since its discovery in the Ivory Coast in 1942, the PPR virus has spread worldwide, reaching China in 2010 and Europe in 2018. Despite the fact that PPR poses a huge threat to the lives of small ruminants and the livelihoods of smallholders, more research is needed into its epidemic potential and transmission speed. Here, we used a combined approach of experimental infections and contact tracing with UWB devices to estimate the probability of transmission of a specific PPRV strain, and to determine the minimum exposure time required for an animal to become infected. Our results indicate that reducing the average distance between animals by a factor of four could reduce exposure time tenfold, indicating a much higher risk for high-density herds compared to roaming ones. Coupled with new estimates of R_0_ and incubation time inferred from data analysis, our work sheds light on the danger posed by PPR. These informations are valuable to veterinarians and policymakers, to better assess the risk of introduction, spread, and impact of PPR, to implement timely and effective interventions.

## Introduction

Peste des petits ruminants (PPR) is a highly infectious and contagious disease affecting small ruminants with morbidity can be as high as 80-90% and a mortality rate as high as 50-80% depending on the epidemiological context. PPR is in the list of economically important animal diseases in which outbreaks must be reported to the World Organization for Animal Health (WOAH). The causative agent of this disease is a virus, the PPR Virus (PPRV). It belongs to the genus Morbillivirus in the family *Paramyxoviridae*. Until 1988, this group of viruses was composed by only four antigenic closely related highly pathogenic viruses, rinderpest virus (RPV) for cattle and buffaloes, measles virus (MV) for human, canine distemper virus (CDV) dog and peste des petits ruminants (PPRV) for sheep and goats [1]. However, since that year, many other viruses have been added to that group: morbilliviruses affecting marine mammals (phocine distemper virus, dolphin morbillivirus, cetacean morbillivirus, feline morbillivirus, porcine morbillivirus and even now a bat morbillivirus) [2], Because RPV and PPRV are affecting ruminants, PPR was for long time ignored in favor of rinderpest. But gene sequence data that had become available as of 1994 onwards have shown that both viruses are different, and that MV and RPV are closer to each other than RPV and PPRV [3]. With the advent of biotechnologies in the 1980s-1990s, tools have been developed for the differential and specific diagnosis of each of those two ruminant diseases. Descripted for the first time in 1942 in Côte d’Ivoire in West Africa, list of countries where the disease is endemic has rapidly expanded as of year 2008 to cover large parts of Africa, from North Africa to Tanzania and Angola at the up-border line of Southern Africa, the Middle and Near East and Asia, nearly all countries from China to Central Asia. Since 2016, this disease is threatening sheep and goat production in Europe [4], with the following countries having experienced some cases: Georgia in 2016 and 2024, Bulgaria from 2018 to 2024, Greece, Romania in 2024, Hungary and Albania in 2025 [5].

The regions that are threatened by PPR are home to about 80% of the world’s sheep and goat population, and the annual losses that are caused by this disease were estimated in 2014 to be at the range of 1.2-2.8 billion (GCES, 2015, [8]). Due to its significant economic impact, particularly on small farmers who rely mainly on sheep and goats for their livelihood in low-income countries, the Food and Agriculture of the United Nations (FAO) and WOAH convened an international conference in 2015 on PPR, conference during which was adopted a strategy for the global eradication of this disease by 2030: the PPR Global Control and Eradication Strategy (PPR-GCES) [6].

PPRV strains that have been identified so far have been grouped into four genetically lineages (I, II, III and IV) based on the virus gene sequence data [7], Initially, this grouping was reflecting the geographical distribution of PPR: lineage I in northern part of West Africa (from Côte d’Ivoire to Senegal), lineage II in the Southern part of West Africa (from Ghana to Nigeria), lineage III in East Africa (Sudan-Ethiopia) and lineage IV in Asia with the Near and the Middle East [7]. However, as of the 2000s, the geographical distribution of PPRV lineages in Africa had started to change with the move of lineage II from its original region to northern parts of West Africa, where lineage I was present, and it has now nearly replaced that latter group. In 2008, PPR broke out in Morocco, the first PPR outbreak in North Africa. The virus strain that was identified in that case was a lineage IV strain [8]. At that period, PPRV strains of lineage IV were identified in Sudan [9], and later in Ethiopia and Eritrea [10,11]. Today, strains of PPRV lineage IV are found in most parts of Africa, either alone or coexisting with viruses of other lineages [8,12–14]. Knowledge of basic parameters of the PPRV transmission from host to host and understanding the virus spread might be useful in describing the transmission dynamics and supporting the PPR eradication program for more efficiency [15]. With that objective in mind, we analyzed data from a set of PPRV experimental infections to estimate basic epidemiological parameters, like contact rate and incubation period of a PPRV strain of lineage IV that was involved in the 2020 PPR outbreaks in Senegal. At the same time, we try to estimate the minimum exposure time, i.e., the minimum time a naïve animal should be in contact with an infected one to get infected. In most cases, in sub-Saharan Africa, the detection and the surveillance of PPR outbreaks are not capillary, and rarely epidemic curves are available that could be used to fit mathematical models. Moreover, serological analysis cannot distinguish between the strains causing the outbreak. This makes it difficult to estimate specific information about the virus. Experimental infection, on the other hand, allows researchers to monitor the temporal behavior of an outbreak in a controlled environment and to estimate parameters by fitting statistical models. Previous works on PPR experimental infection have focused on the pathogenesis of PPRV [16,17]. However, the study by Herzog et al. (2020 and 2024) used data from experimental infection in cattle to estimate the transmission rate, R_0_, in this species and its eventual role in PPRV transmission [18–21].

Several factors could impact this measure, one being the virulence/transmissibility of the virus, the status of the animals, and the contact pattern among them. PPRV, like other morbilliviruses, spreads mainly through direct contact between susceptible hosts and those excreting the virus. Along normative days, the contact patterns among animals change due to their relative movements and circadian activities and could impact the probability of virus transmission from an excreting animal to a naïve one. UWB and Radio-frequency identification (RFID) devices are wearable devices that have been widely used to study interaction among individuals. Adapted to work in confined environments, this type of device provides more detailed and frequent information than GPS. The signals of the two devices are frequently triangulated, and the distance between their bearers is estimated at each time step. The result is a dynamic contact network among individuals. UWB and RFID devices have been widely used to study interactions among individuals (humans) in different settings like social events, conferences [22,23], hospitals [24], schools [25,26], and villages [23,27,28]. The results provide information for studying the interactions among individuals and retrieving information about possible drivers. Similarly, UWB devices have been recently applied to study animal behavior and welfare for different animal species, dogs, chickens, sheep, cows, and horses [29–37]

In our study, we analyzed data from experimental infections where animals were equipped with UWB devices to collect information about contacts, and a set of statistical models was developed to infer the value of the virus’s transmissibility. This article is divided in two parts : in the first one, we estimated epidemiological parameters of interest, like transmission rate R_0_ and incubation time without considering the impact of interaction patterns; in the second part we develop a suite of Bayesian statistical models to estimate the probability of transmission and consequently the minimum time of exposition for an animal to get infected. To our knowledge, this is the first time that this kind of study (combining infection monitoring and contact patterns) has been performed. By taking account of contact variation, the analyses could help provide more accurate estimates of transmission probabilities and, consequently, the impact of PPR.

## Results

### Observed contact distribution and patterns

In our analysis, we considered only those experiments for which we had a complete dataset covering the entire duration of the experiment.

Fig 1A shows the distance distribution among animals for all experiments. From a visual inspection, the distance distribution appears different between experiments, which is confirmed by the Kruskal-Wallis test results (*χ*^2^= 1331181, p-value < 2.2e-16) indicating that some of the distributions are different, and the *post-hoc* pairwise Wilcox test, indicating that there is a significant difference among all the experiments (all p-value < 2.2e-16) except two (namely the experiments 1_2 and 2_2) for which the corresponding p-value = 0.19.

**Fig 1.**
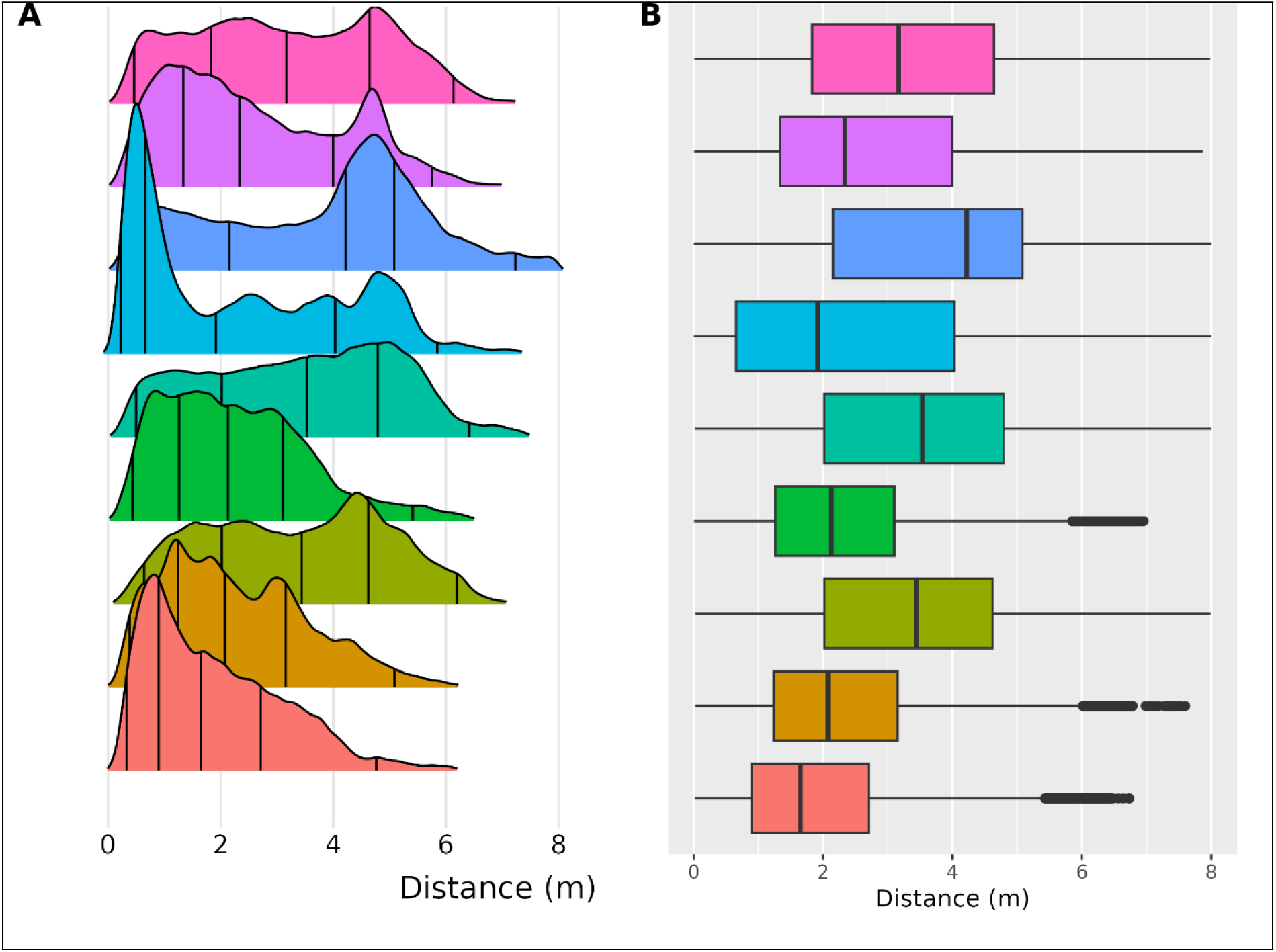
**(A) Distribution of distances among animals per experiment.** Colors correspond to different experiments. Vertical lines indicate quartile values of the distribution. **(B) Boxplot to resume the distribution.**

Fig 1B summarizes the distribution in the form of a boxplot. As we can notice, the median distance in each experiment varies. In particular, we notice that for five of them the median is in the range 1.5-2.5 m., while for the others it is between 3.3 and 4.5 m. The differences among contact patterns could indicate different behavior of the animals, due to external reasons, and could impact the transmission of the virus. We analyzed the temporal network to identify possible eras, i.e., specific periods related to different collective behaviors that could indicate the presence of circadian activity. In Fig S5 in S1 File, the median (solid line) and the minimum and maximum distances among animals are reported for each second of the exposure phase. The era identification analysis shows inconclusive results: a single large period was identified, lasting more than 90% of the time, together with some very short eras (of the order of the minutes).

Instead of focusing on the characteristics of the overall network, we extracted information about the contact pattern between the seeder and each susceptible animal (called proximity analysis). The proximity results presented in Fig 2 shows the fraction of time susceptible animals spent in very close contact(less than the 0.5 m), medium range contact (0.5-1.0 m), long range (1;0-2.0 m) and distant contact (more than 2.0 m) The pattern of dynamic behavior is characterized by frequent, spontaneous fluctuations, where susceptible individuals intermittently reduce their proximity to the seeder before increasing the distance again. Specifically, susceptible individuals would briefly enter closer proximity zones (under 1 meter) before returning to the mostly distancing pattern of beyond 2 meters. Overall, individuals spend a very small fraction of time in close contact with the seeder, while most of the time they are at a large distance.

**Fig 2.**
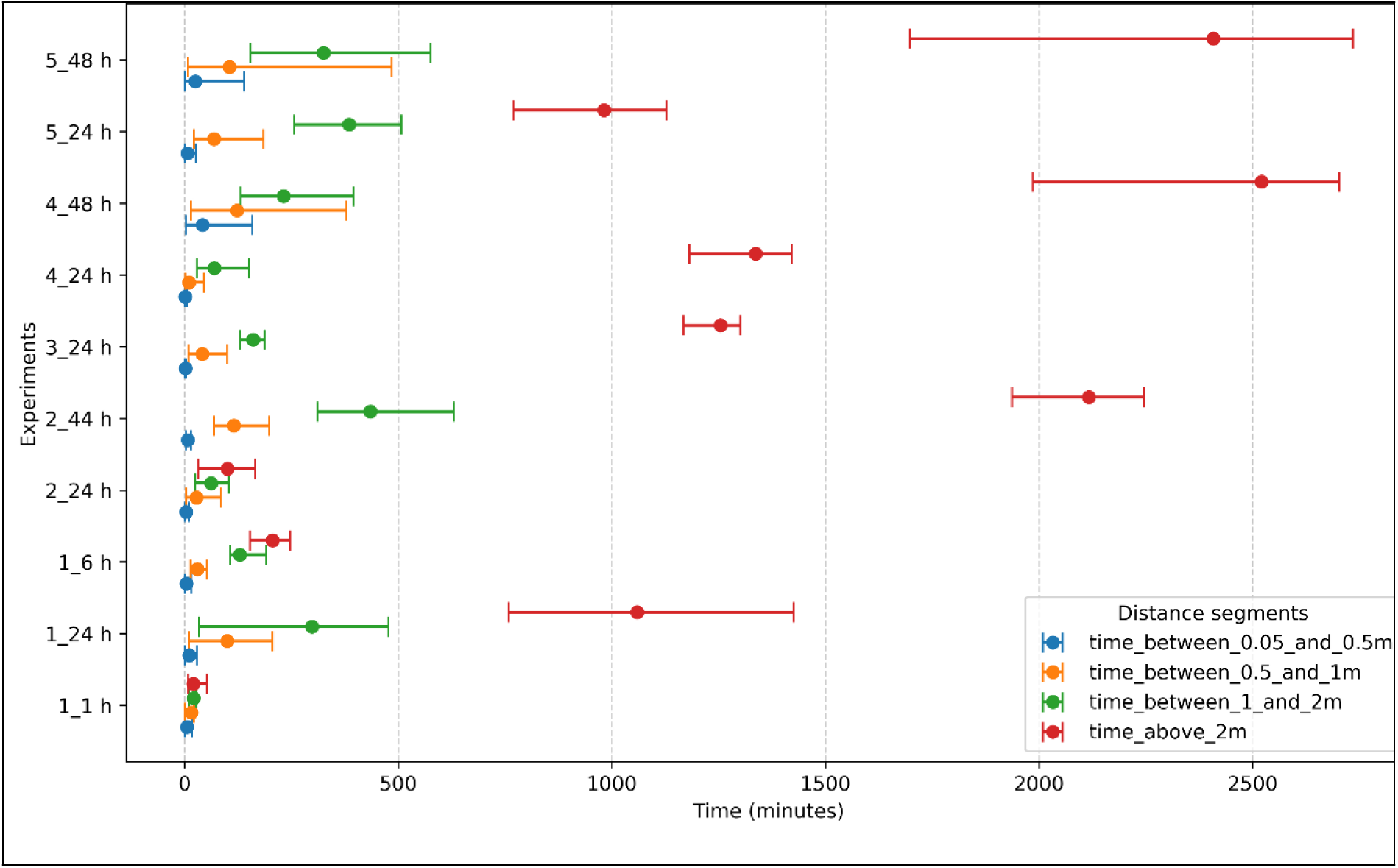
Contact time between seeders and susceptible animals’ distribution under distance segments. For each experiment, the color of each bar corresponds to a specific range of distance, the dot corresponds to the average time a susceptible animal is at a certain distance from the seeder one, and the extreme indicates the minimum and maximum time.

### Clinical Observations and Serology results

In this section, we present the results of the tests and observations conducted to assess the animals’ health status and to determine whether they were infected (through RT-PCR), as well as their immune response to the exposure to the pathogen (through serological testing). The data collected from each experiment, including the monitoring period and the number of positive serological results, are summarized in Table 1. Overall, 18 positive cases based on positive serology were identified out of 70 susceptible individuals. A significant association between exposure time and infection status was found (Fisher Exact test, p-value < 0.001), indicating that exposure duration influences infection likelihood. Three of these (ID 32L, 42L and 81) were found to be positive during post-mortem analysis, while the others exhibited an immune response (positive serology) during the monitoring period. Only five animals (ID 104, 108, 99, 4 and 4D) were also positive in the RT-PCR test, and of these, only three (ID 104, 108 and 99) exhibited symptoms such as nasal and ocular discharge, diarrhea (Table 2).

**Table 1.** Summary of experimental infection results by experiment session and duration.

| Experiment | Duration | Date | Batch size<br>(In-contact animals) | Positive |
| --- | --- | --- | --- | --- |
| 1 | 1 h | 27/06/2021 | 6 | 0 |
|  | 6 h | 27/06/2021 | 6 | 0 |
|  | 24 h | 27/06/2021 | 6 | 0 |
| 2 | 24 h | 25/08/2021 | 6 | 0 |
|  | 24 h | 26/08/2021 | 6 | 1 |
|  | 44 h | 27/08/2021 | 6 | 4 |
| 3 | 24 h | 02/03/2022 | 5 | 1 |
|  | 24 h | 02/03/2022 | 5 | 3 |
| 4 | 24 h | 28/04/2022 | 6 | 0 |
|  | 48 h | 29/04/2022 | 6 | 4 |
| 5 | 24 h | 14/06/2022 | 6 | 1 |
|  | 48 h | 15/06/2022 | 6 | 4 |

We report the number of newly infected animals (Positive) after the exposure phase.

**Table 2.** Day of symptom onset, RT-PCR positivity, serology positivity, and death for the confirmed cases of PPR by experiment and ID, as the number of days elapsed since the end of the exposure phase.

| Session | Batch | Starting date | Contact duration | Animal Id | Symptoms | RT-PCR | Serology | Death |
| --- | --- | --- | --- | --- | --- | --- | --- | --- |
| 2 | 2 | 26/08/2021 | 24 h | 40 |  |  | 20/09/2021 |  |
|  | 3 | 27/08/2021 | 44 h | 104 | 04/09/2021 | 06/09/2021 | 10/09/2021 | 13/09/2021 |
|  |  | 27/08/2021 | 44 h | 108 | 08/09/2021 | 08/09/2021 | 12/09/2021 |  |
|  |  | 27/08/2021 | 44 h | 128 |  |  | 20/09/2021 |  |
|  |  | 27/08/2021 | 44 h | 99 | 04/09/2021 | 07/09/2021 | 06/09/2021 | 07/09/2021 |
| 3 | 1 | 01/03/2022 | 24 h | 20 |  |  | 15/03/2022 | 18/03/2022 |
|  | 2 | 02/03/2022 | 24 h | 32L |  |  |  | 14/03/2022 |
|  |  | 02/03/2022 | 24 h | 42L |  |  |  | 13/03/2022 |
|  |  | 02/03/2022 | 24 h | 23KD |  |  | 16/03/2022 | 20/03/2022 |
| 4 | 2 | 29/04/2022 | 48 h | 4 |  | 18/05/2022 |  | 18/05/2022 |
|  |  | 29/04/2022 | 48 h | 4D |  | 16/05/2022 |  | 16/05/2022 |
|  |  | 29/04/2022 | 48 h | 16 |  |  | 15/05/2022 |  |
|  |  | 29/04/2022 | 48 h | 41L |  |  | 07/06/2022 |  |
| 5 | 1 | 14/06/2022 | 24 h | 6 |  |  | 06/07/2022 |  |
|  | 2 | 15/06/2022 | 48 h | 18 |  |  | 24/06/2022 |  |
|  |  | 15/06/2022 | 48 h | 33 |  |  | 08/07/2022 |  |
|  |  | 15/06/2022 | 48 h | 59 |  |  | 24/06/2022 |  |
|  |  | 15/06/2022 | 48 h | 81 |  |  |  |  |

Symptoms indicate the day when any of the symptoms (ocular, nasal discharge, diarrhea, high temperature) were recorded, virology the day when samples were positive to the lab test, serology the day when antibodies were detected in the blood, and death the day animals were found dead and positive to the disease.

Fig 3 visually depicts the proportions of infected versus non-infected individuals across different exposure time categories, emphasizing that no infections were recorded for exposure times under 24 hours. We observed that experiments with longer durations of exposure resulted in a higher number of infections, indicating that the length of the exposure period may impact the probability of transmission.

**Fig 3.**
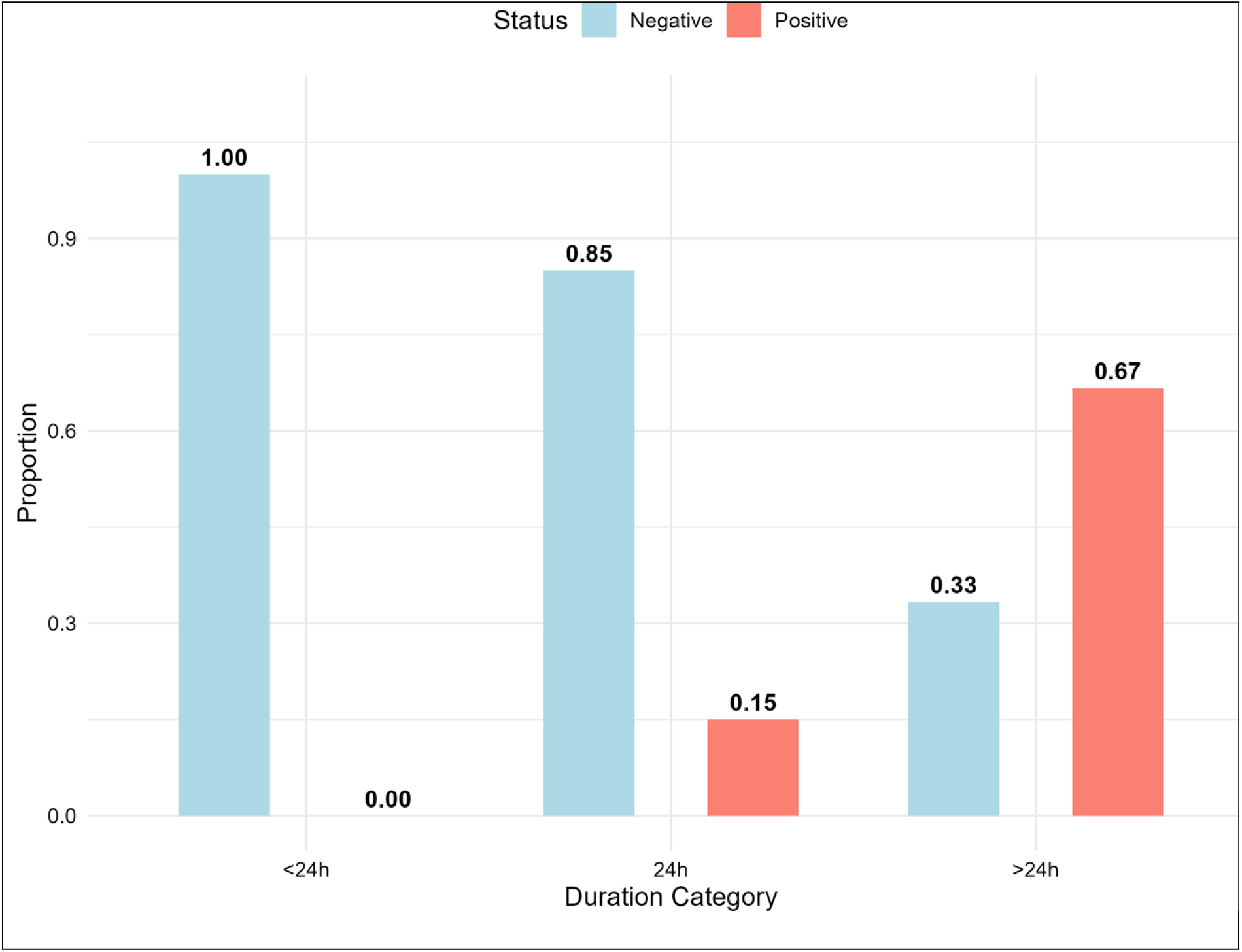
Proportions of infection by exposure time categories. Outputs of the experimental infections are grouped based on the duration of the exposure phase. Red corresponds to the fraction of positive in each category, Cyan to the negative ones.

### Transmission rate estimation

The transmission rates were initially estimated on an hourly basis. However, for reporting purposes, these values were multiplied by 24 to convert them into daily rates. The estimated value of *β* is 0.86 (95% highest density interval (HDI): 0.62 – 0.94).

The basic reproductive number R_0_ is estimated to be 4.3 (95% HDI: 3.1 – 4.8) when considering an infectious period of 5 days as used by El Arbi et al. [38] and 8.6 (95% HDI: 6.2 – 9.6) when considering an infectious period of 10 days, as reported by Fournié et al. (2018) and Herzog et al. (2024).

### Incubation period estimation

Table 2 shows, for each positive confirmed animal, the day at which it showed symptoms (symptoms), was positive to RT-PCR or serological tests, and in case of deaths, the day of death. For the survival analysis, we considered the first day on which one of these factors was registered. As the table shows, very few positive animals (only 3) showed symptoms during the follow-up period, indicating that the virus could circulate silently in a herd.

Preliminary survival analysis was done to check if there was a difference between exposure duration and length of the incubation period. The log-rank test showed no significant difference between the groups (Chi-square 0.1, p-value 0.8). Because of this, we conducted survival analysis without taking into consideration the duration of the exposure phase. Among the three parametric models, the LogNormal one outperformed the others (Table 3). The average incubation period is around 14.50 days (95% CI 11.8-20.2).

**Table 3.** Estimation of the incubation period using three different parametric distributions (Weibull, Gamma, and LogNormal) showing the median of the parameter estimates, the 95% Credible Interval, Leave-one-out cross-validation information criterion (LooIC) and Watanabe Akaike Information Criterion (WAIC)

|  | **2.5%** | **50%** | **97.5%** | **WAIC** |
| --- | --- | --- | --- | --- |
| **Weibull** | 11.8 | 15.4 | 20.2 | 119.5 |
| **Gamma** | 11.7 | 14.6 | 18.5 | 117.9 |
| **LogNormal** | 11.6 | 15.2 | 22 | 116.6 |

Fig 4 shows the empirical Kaplan-Meier curve, and the theoretical one obtained using a lognormal distribution. During the first four days after exposure, none of the animals can secret the virus. From the fifth day till the 20^th^ day, the probability of not developing symptoms decreases. Around the 15^th^ day, half of the population is no longer incubating, and animals either show symptoms or are tested positive. After the 22^nd^ day, only a small percentage of animals still develop symptoms.

**Fig 4.**
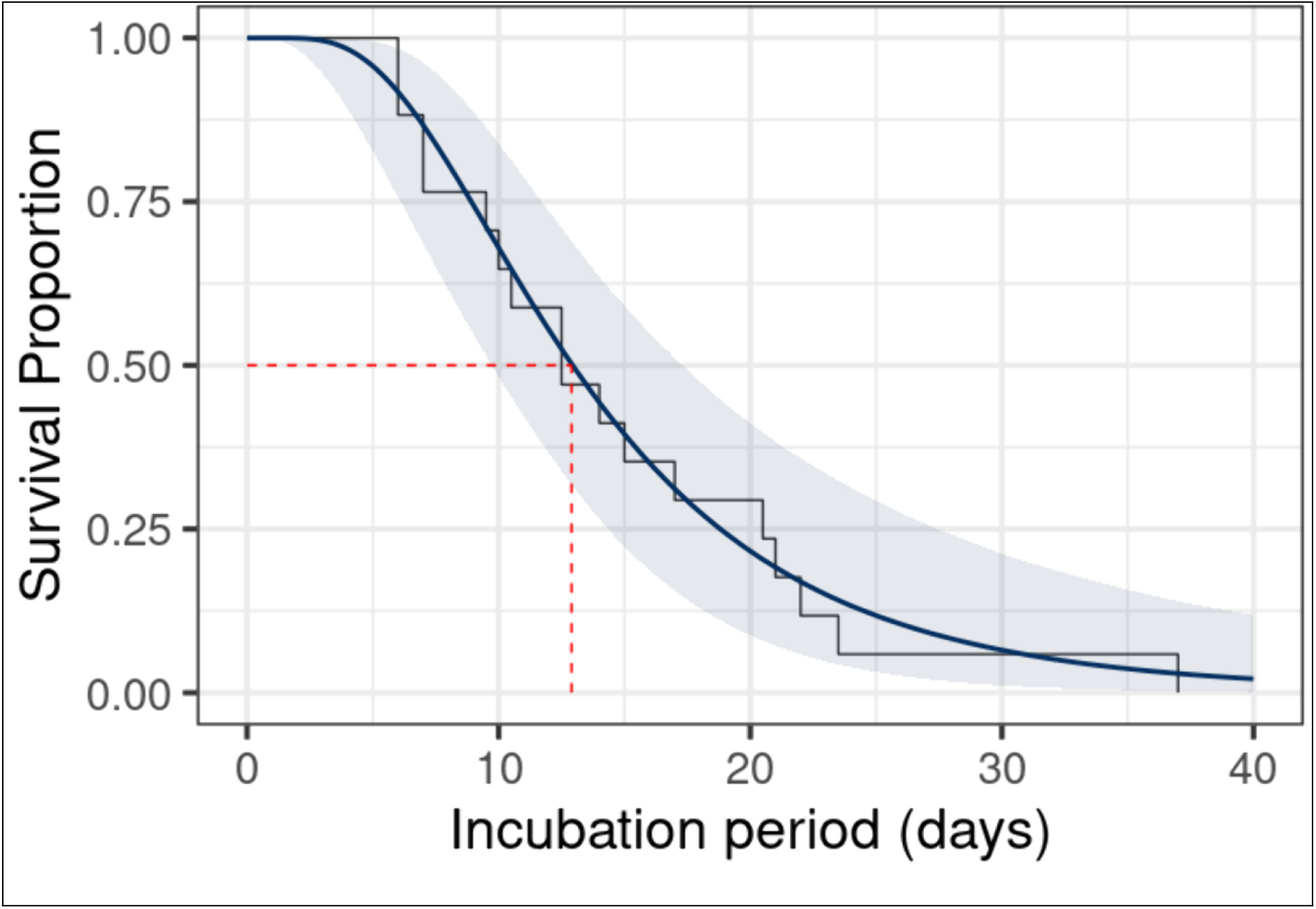
Kaplan-Meier curve for the empirical (black solid line) and the estimated one (blue). The shaded area corresponds to the 95% C.I. of the survival plot. The dashed line indicates the time (in days) half of the population has become infectious.

### Transmission probability estimates

In Table 4, an overview of the prior distributions used for each parameter across all models is presented. For the transmission probability parameters, we employed Beta distributions to constrain values between 0 and 1, reflecting the probabilistic nature of these parameters. The decay parameter in the envelope model was assigned a Gamma distribution to ensure positivity while allowing for a range of plausible values. For scaling factors in the Stratified model, we used a uniform prior over a range that encompassed biologically reasonable values.

**Table 4.**
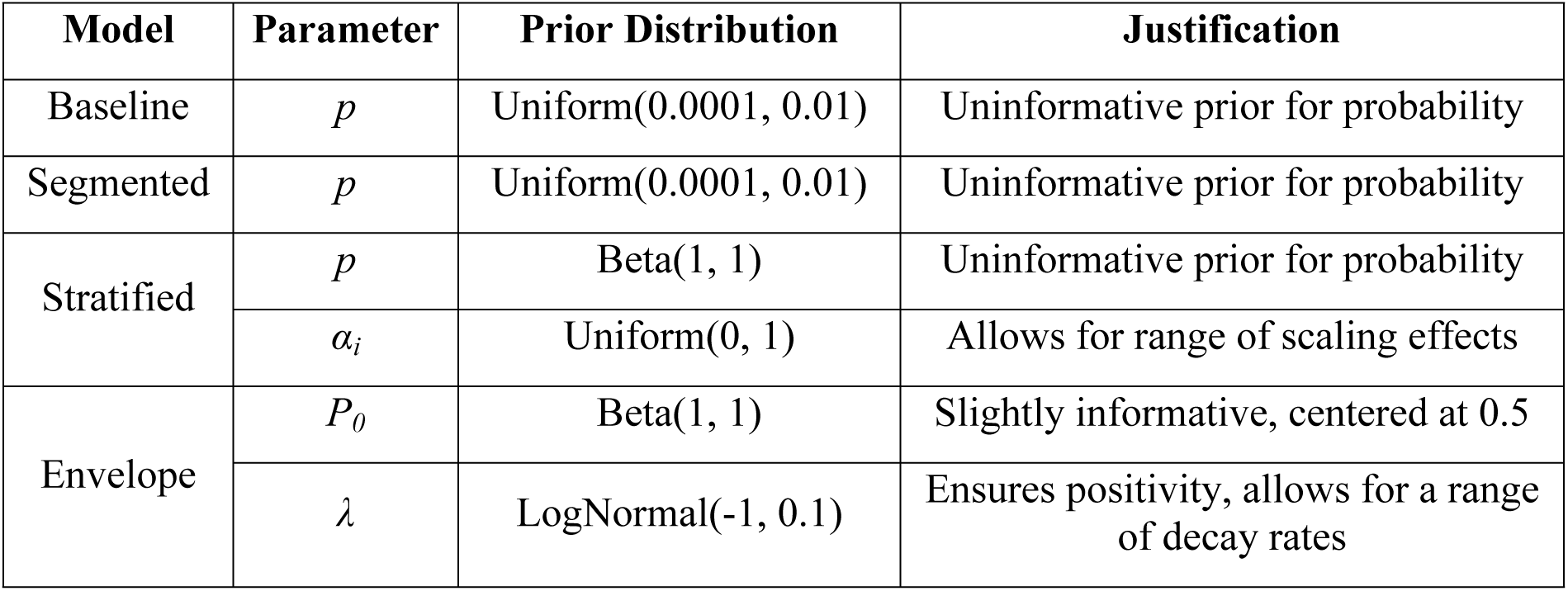
Prior distributions for model parameters.

The estimated probability parameters *p* from MCMC, differ across models depending on how they incorporate distance. The results of the inference through calibration are shown in Table 5. Probability estimates for the Baseline case are the lowest. This is mainly due to the fact that in the model, individuals are considered always to be exposed independently of the distance.

**Table 5.** Summary of posterior distributions for all models.

| Model | Parameter | Mean | SD | HDI 95% lower | HDI 95% upper |
| --- | --- | --- | --- | --- | --- |
| Baseline | $p$ | 0.0003 | 0.0001 | 0.0001 | 0.0004 |
| Segmented 1 m | $p$ | 0.0063 | 0.0014 | 0.0037 | 0.0092 |
| Segmented 1.5 m | $p$ | 0.0028 | 0.0007 | 0.0015 | 0.0041 |
| Segmented 2 m | $p$ | 0.0014 | 0.0003 | 0.0008 | 0.0021 |
| Envelope 1 m | $P_0$ | 0.0112 | 0.0029 | 0.0059 | 0.0169 |
| | $\lambda$ | 0.3702 | 0.0374 | 0.297 | 0.4428 |
| Envelope 1.5 m | $P_0$ | 0.0043 | 0.0011 | 0.0023 | 0.0063 |
| | $\lambda$ | 0.3703 | 0.0371 | 0.2999 | 0.4439 |
| Envelope 2 m | $P_0$ | 0.0025 | 0.0006 | 0.0014 | 0.0038 |
| | $\lambda$ | 0.3706 | 0.0372 | 0.2999 | 0.445 |
| Stratified 1 | $p$ | 0.0046 | 0.0043 | 0.0003 | 0.0133 |
| | $\alpha_1$ | 0.3764 | 0.2708 | 0.014 | 0.9012 |
| | $\alpha_2$ | 0.5064 | 0.2627 | 0.1092 | 0.9973 |
| | $\alpha_3$ | 0.5879 | 0.2524 | 0.1586 | 0.9999 |
| Stratified 2 | $p$ | 0.0117 | 0.0063 | 0.0036 | 0.0245 |
| | $\alpha_1$ | 0.5855 | 0.2427 | 0.1835 | 0.9996 |
| Stratified 3 | $p$ | 0.0006 | 0.0003 | 0.0002 | 0.0014 |
| | $\alpha_1$ | 0.4917 | 0.2644 | 0.0869 | 0.9895 |
| Stratified 4 | $p$ | 0.0016 | 0.0011 | 0.0002 | 0.0038 |
| | $\alpha_1$ | 0.41 | 0.266 | 0.0264 | 0.9192 |
| | $\alpha_2$ | 0.5531 | 0.2603 | 0.1281 | 1 |
| Stratified 5 | $p$ | 0.0064 | 0.0049 | 0.0011 | 0.0163 |
| | $\alpha_1$ | 0.4705 | 0.2621 | 0.061 | 0.9486 |
| | $\alpha_2$ | 0.5866 | 0.2476 | 0.1693 | 1 |

For the other models, spatial information was incorporated *post hoc,* by applying distance-based filters to the proximity data after it was collected; no constraints were imposed during data generation or experimentation. Segmented models used simple distance filters (e.g., allowing transmission only within 1 m, 1.5 m, or 2 m) produced progressively lower *p* estimates (*p* = 0.0063, 0.0028, and 0.0014, respectively), showing that increasing the spatial window reduces the estimated per-individual transmission probability. Stratified models further refined this approach by dividing the spatial range into intervals (e.g., 0.05-0.5 m, 0.5-1 m, 1-1.5 m, etc.), and we noticed that the more models cover close-contact (0.05-0.5 m) the more the probability *p* estimates increase (e.g., Stratified 2, Stratified 5 and Stratified 1), and for the case when models integrated distances above 2 m, the *p* estimates decrease (e.g. Stratified 3). In particular, we noticed that for Stratified 2 model the probability of transmission at very close distance (less than 0.5 m) is 39 times larger than in the baseline case. This result indicates that spatial density strongly impacts the diffusion of pathogens.

### Model evaluation results

The ROC curve analysis was performed to assess the diagnostic performance of our models, with the mean AUC and its 95% HDI presented in Table 6. Fig 5 illustrates the average ROC curves for these models, showcasing their sensitivity and specificity across various thresholds. The analysis of AUC values provided critical insights into the predictive capabilities of the evaluated models. Notably, the baseline model and the Stratified 4 model exhibited the highest AUC values, indicating superior predictive accuracy, with the baseline model achieving an AUC of 0.84 and the Stratified 4 model closely following at 0.82. These findings suggest that both models are highly effective in distinguishing between different classes. Additionally, the Stratified 3 and Stratified 1 models demonstrated similar AUC values of 0.81 each, reflecting similar levels of predictive efficiency. In contrast, the segmented and envelope models yielded lower AUC values overall; however, within their respective categories, the segmented model and the envelope model performed relatively better at distances below 2 meters, achieving AUC values of 0.73 and 0.72, respectively. These results underscore the robustness of both basic and Stratified models in predictive tasks while highlighting potential limitations associated with segmented and envelope models.

**Fig 5.**
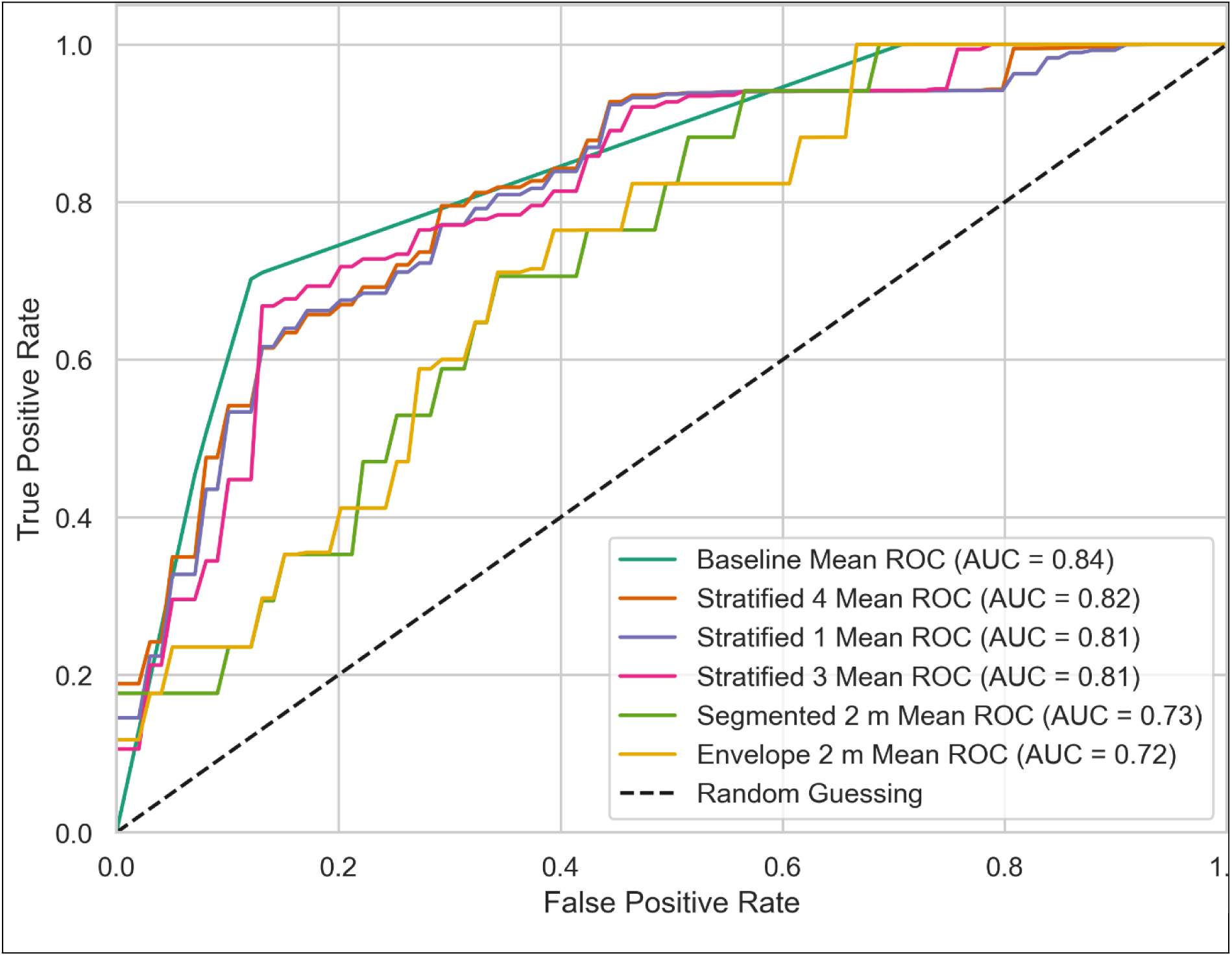
Mean ROC curves of best models. Colors correspond to the best models identified in each category and compared to the baseline case when no contact information is considered. The average ROC and AUC are estimated from the average over a parameter sample of 100 000 transmission probability values.

**Table 6.** Comparison of AUCs for different models.

| Model | Mean AUC | SD | HDI 95% lower | HDI 95% upper |
| --- | --- | --- | --- | --- |
| Baseline | 0.84 | 0 | 0.84 | 0.84 |
| Segmented 1 m | 0.67 | 0 | 0.67 | 0.67 |
| Segmented 1.5 m | 0.68 | 0 | 0.68 | 0.68 |
| Segmented 2 m | 0.73 | 0 | 0.73 | 0.73 |
| Envelope 1 m | 0.65 | 0 | 0.65 | 0.65 |
| Envelope 1.5 m | 0.68 | 0 | 0.68 | 0.68 |
| Envelope 2 m | 0.72 | 0 | 0.72 | 0.72 |
| Stratified 1 | 0.81 | 0.02 | 0.76 | 0.84 |
| Stratified 2 | 0.66 | 0.01 | 0.64 | 0.67 |
| Stratified 3 | 0.81 | 0.01 | 0.78 | 0.83 |
| Stratified 4 | 0.82 | 0.02 | 0.76 | 0.84 |
| Stratified 5 | 0.71 | 0.02 | 0.68 | 0.73 |

In Table 7, we summarized the performance of the models based on the WAIC results. In the table “elpd_waic”: This is the expected log pointwise predictive density (ELPD). High values indicate better predictive accuracy. “p_waic”: This is the effective number of parameters in the model that provides an estimate of model complexity; “elpd_diff”: This is the difference in elpd_waic between the current model and the best model. Smaller differences indicate that the models have similar predictive performance; “se”: The standard error of ELPD estimate. “rank” indicates the ranking of models based on their WAIC values. A low rank indicates a better model.

**Table 7.** WAIC comparison of the different models to assess their performance and complexity.

| Model | elpd_waic | p_waic | elpd_diff | se | dse | rank |
| --- | --- | --- | --- | --- | --- | --- |
| Baseline | -27.93 | 0.88 | 0 | 2.45 | 0 | 1 |
| Segmented 1 m | -35.55 | 1.07 | 7.62 | 5.52 | 4.49 | 10 |
| Segmented 1.5 m | -35.28 | 1.12 | 6.99 | 5.41 | 4.5 | 8 |
| Segmented 2 m | -31.52 | 1.04 | 3.59 | 3.75 | 2.61 | 5 |
| Envelope 1 m | -35.57 | 1.27 | 7.64 | 6.41 | 4.99 | 11 |
| Envelope 1.5 m | -35.28 | 1.12 | 7.36 | 5.61 | 4.68 | 9 |
| Envelope 2 m | -32.11 | 1.07 | 4.18 | 4.05 | 2.97 | 6 |
| Stratified 1 | -28.95 | 1.25 | 1.02 | 3.39 | 1.9 | 4 |
| Stratified 2 | -36.31 | 1.32 | 8.39 | 5.58 | 4.53 | 12 |
| Stratified 3 | -28.77 | 1.15 | 0.84 | 3.29 | 1.76 | 2 |
| Stratified 4 | -28.88 | 1.30 | 0.95 | 3.32 | 1.86 | 3 |
| Stratified 5 | -32.29 | 1.30 | 4.37 | 4.02 | 2.87 | 7 |

We compared multiple models using WAIC. The baseline model had the highest ELPD (elpd_waic = –27.93) and thus the best out-of-sample predictive performance. The Stratified 3, Stratified 4, and Stratified 1 models showed minor decreases in ELPD relative to the baseline (elpd_diff = 0.84, 0.95, and 1.02 respectively), with standard errors that suggest these differences are not substantial (dse = 1.76, 1.86, and 1.9 respectively). All of these models had similar complexity, with p_waic values between 0.88 and 1.3.

The remaining models Segmented, Envelope and some Stratified models had notably lower ELPD indicating some problems in predictive performance. The elpd_diff values for these models were larger than the associated standard errors (se), supporting the conclusion that these models predict less well than the baseline and top Stratified models.

### Goodness of fit

We conducted Bernoulli simulations using the estimated probability values *p* derived from the best-ranked models: Baseline, Stratified 4, Stratified 3, and Stratified 1, to generate binary outcomes (0 or 1) for each scenario. These simulations were grouped by experiments, and the total number of positive cases per experiment was calculated by summing the binary outcomes. The results are summarized in Fig 6, which compares the predicted mode number of positive cases against the real observed values. The results demonstrate that the four models are nearly juxtaposed, indicating that their performance in terms of fitting and prediction is almost identical. This suggests a high level of consistency across the models in capturing the underlying patterns of the data. Additionally, the predicted curves closely follow the trends of the real observations, further validating the models’ accuracy. However, an exception is noted in experiment 2_24, where the predicted outcomes deviate slightly from the observed trends.

**Fig 6.**
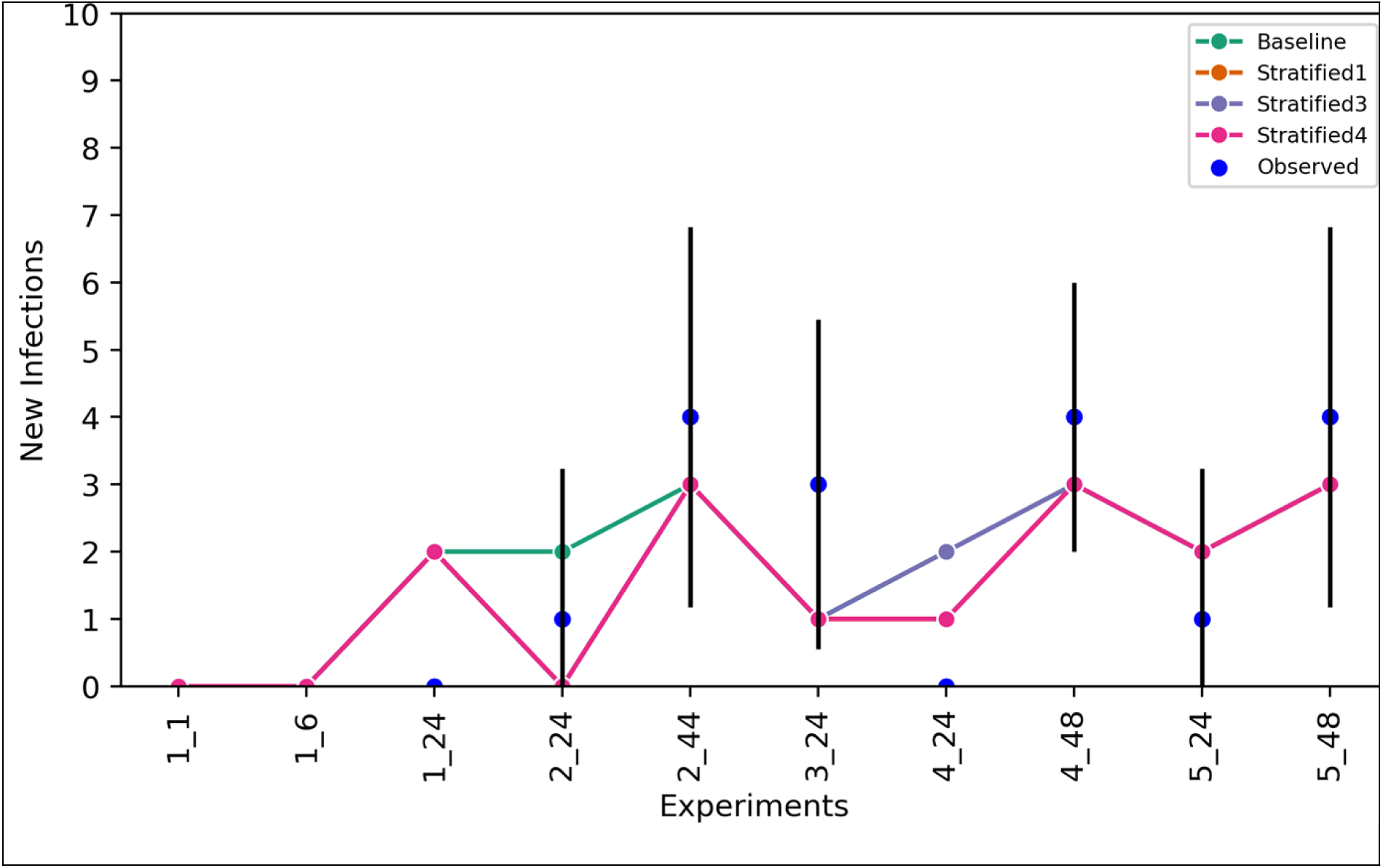
Comparison of model-predicted and experimentally observed PPR positive cases. Predictions are displayed as mode values, with error bars indicating the standard deviation, and real observations are presented as blue dots.

### Exposure time estimation

To estimate the minimum exposure time, we simulated various exposure scenarios. In Fig 7 we report the results of the estimation of the probability for an individual to get infected while in contact with an infected animal after a certain amount of time. We considered several scenarios, based on assumptions about contact patterns. We initially used the estimated values of the probability of success from the baseline to determine the minimum exposure time, measured in hours, needed for successful contamination. In this example, we assumed that animals could move freely and that the chance of infection is uniform. If one of them is infected, there is a 50% chance for a naïve animal to catch the infection after being exposed for about 38.50 hours corresponding to one day and a half. On the other hand, if the probability of infection is 0.95, it is estimated that the average exposure time is around 166.40 hours (almost one week). These estimations are based on the fact that animals can move freely and interact at short distances rarely. However, livestock markets are very crowded places, where thousands of animals spend 6-8 hours in very close contact. To estimate the exposure times in these conditions, we considered simulations using the parameters for close contacts estimated by our models.

**Fig 7.**
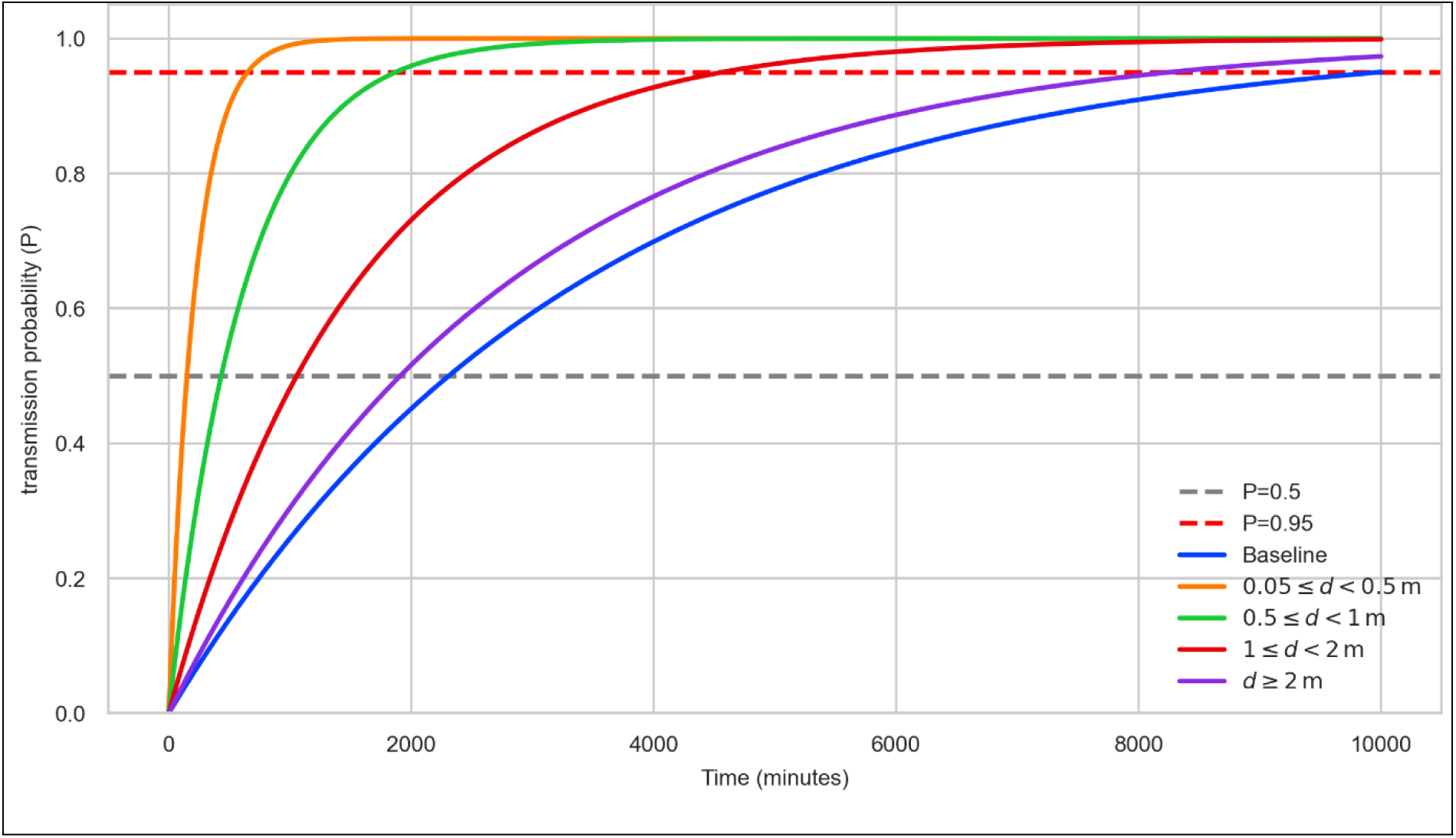
Estimating the minimal exposure duration required to achieve contamination. The solid lines represent the cumulative probability of an animal becoming infected after being exposed for ***t*** minutes. Colors indicate different scenarios: the baseline case without contact patterns, and hypothetical cases where animals remain at fixed distances, Stratified 1 was used for the first distance segment 0.05-0.5 m, and Stratified 4 for the remaining segments. Dashed lines mark the thresholds corresponding to 50% and 90% infection probability.

The analysis of simulation using the Stratified 1 and Stratified 4 models revealed a consistent positive relationship between exposure time and the inter-individual distance. We used Stratified 1 model to estimate the exposure time related to a fixed distance within the first range of 0.05–0.5 m and Stratified 4 for the rest of distance segments. The exposure duration increased with distance to reach the probability thresholds of 0.5 and 0.95. To reach a transmission probability of 0.5, the mean exposure durations were 2.51 hours, 7.21 hours, 17.60 hours, and 31.83 hours for the distance segments 0.05–0.5 m, 0.5–1 m, 1–2 m, and ≥2 m, respectively. To reach a probability of 0.95, the required durations increased substantially, with corresponding values of 10.83 hours, 31.18 hours, 76.09 hours, and 137.58 hours, respectively, across the same distance intervals. These estimates indicate a substantial increase in the average exposure time required to achieve higher probabilities of infection, highlighting the variability and escalation of exposure duration necessary as the risk of infection increases.

## Discussion

The transmissibility and other epidemiological parameters of PPRV are difficult to estimate from field data. Most mathematical models that were developed up to date rely on serological data [21,38,39]. Therefore, there was a need to undertake experimental infections to closely monitor the evolution of the virus spread and estimate the epidemiological parameters of interest. In this work, we have investigated PPRV transmission from goat to goat by using a Senegalese PPRV strain of lineage IV [40] by analyzing data from an experimental infection. Our main interests were to estimate the transmission rate and incubation time, estimate the minimum exposure time needed for an animal to become infected and understand how the contact pattern among animals could impact the transmission of the virus.

Our experimental results yielded an estimated daily transmission rate *β* = 0.86 (95% HDI: 0.62 – 0.95) which at its turn corresponds to a value of R_0_ of 4.3 (95% HDI: 3.1 – 4.8) when considering an infectious period of 5 days as used by ElArbi et al. (2019), and 8.6 (95% HDI: 6.2 – 9.6) when considering an infectious period of 10 days as reported by Fournié et al. (2018), and Herzog et al. (2024). Previous estimates of both parameters vary depending on the type of study (modeling or experimental infection), geographical context (Western or Eastern Africa), husbandry practices (pastoralist against sedentary) as well as the value of the infectious period considered. In the context of experimental infection, Herzog et al. (2024) have reported a transmission rate from small ruminants to small ruminants of 0.26 (95% HDI: 0.14 – 0.63), which is approximately more than three times lower than ours, while in modeling studies in Mauritania is around 1.61 [38], and in Ethiopia between 0.15 and 0.62 [39], depending on the husbandry practice. Similarly, the estimates for R_0_ widely vary from around 1.5 in highlands Ethiopia (mostly sedentary agropastoral) [39], to 2.8 and 2.9 in Senegal and Mauritania [40, 37], till 4 and 6 in Tanzania [42] and lowland Ethiopia Besides the aforementioned factors, explanation for these differences can be trace back in experimental protocol. In our study, we considered only young goats who were kept strictly separated and isolated at the end of each exposure phase. This last precaution has been taken to ensure that transmission events were limited to controlled periods of exposure. In the abovementioned studies, herds were mixed (either goat and sheep, either small and large ruminants) and animals of different ages were mixed. Because of this, animals with different susceptibilities to the disease were mixed thus reducing the number of effective contacts. Furthermore, in the abovementioned studies, parameters were estimated from final sizes and serology, while in our study the duration of the experiment was fixed a priori, and the focus was more on the number of animals that could be infected in a time window. In the study by Herzog et al. (2024), the animals were kept in common pens throughout the infectious period of the experiment, which enabled continuous contact between infected and susceptible animals, with possibilities of secondary infections in that case, and the statistical model was calibrated on data collected over several days. Because of this the number of new infections could decrease over time and the transmission parameter could represent a sort of temporal average. In our study, instead, contact between infectious and naïve animal for a fixed period of time in view of avoiding secondary sources of infection.

Until now, the average incubation period of PPR is considered to range from 4 to 6 days, though it can range from 3 to 10 days. For surveillance and control purposes, the WOAH recommends adopting a conservative incubation period of 21 days [43]. In this study, our data showed a substantially longer incubation period, around 15 days (95% CI: 12.55–23.28). This prolonged latency, combined with the fact that only a small proportion of infected animals exhibit clinical symptoms, represents a significant challenge for effective disease detection and containment. In most parts of Africa, where animals are kept mobile for different reasons such as transhumance, a long incubation period could mean that asymptomatic animals could travel several hundred km, and cross several national borders, before disease could be detected. This poses a threat to the control of disease and demands stricter and more harmonized surveillance measures among countries in the area.

Direct transmission requires close physical contact between animals. The concepts of “contact” and “effective contact” are fundamental yet complex ideas in mathematical epidemiology. For contact to be effective, animals must be close enough for droplets to travel from infected to susceptible individuals. However, throughout a typical day, animals move and interact at different times and for varying durations. As a result, the possibility of pathogen transmission is shaped by the pattern of effective contacts. To monitor activity patterns and distances between animals, high-resolution Ultra-wideband (UWB) sensors have proved to be powerful tools.

Previous studies using networks constructed from data collected using UWB, or similar, devices employed threshold models to identify effective contacts (when two individuals were within 1– 1.5 meters of each other) (http://www.sociopatterns.org/) based on a literature review [44–49]. For PPR, there is limited, if not none, information about transmission distances that could inform our model. Therefore, we calibrated statistical models with multiple distance-based hypotheses to account for activity patterns and heterogeneity, and to evaluate how proximity affects pathogen transmission. Our results indicate that individuals spend a very short amount of time—just a few minutes—close to each other. Moreover, the analysis of the temporal contact network revealed no clear contact patterns, suggesting that animals tend to avoid close interactions. This behavior may be due to a lack of familiarity, the animals in our study were young and, coming from different herds, and were not previously acquainted. This could have influenced their social behavior and reduced their gregariousness, especially considering that young animals typically remain close to their mothers and siblings. We are currently analyzing the footage by cameras that were in the animal facility. The results, to be reported in a second manuscript, will give more information about the contacts between animals in the same premises.

Despite ongoing research, there is little information on the minimum exposure period required for successful contamination. To this end, it becomes important to estimate a probability of transmission for “effective contact”. In the absence of a definition of close contact for PPR transmission, several models have been developed that considered different definitions of effective distance and transmission probabilities that decrease with distances. Nevertheless, the baseline model, that did not incorporate any information about contact patterns, slightly outperformed all other models based on the WAIC and the AUC-ROC. The observation that the simplest model, which does not incorporate distance, provided the best fit suggests that certain aspects of our experimental design or initial assumptions about PPRV transmission dynamics may need to be reconsidered.

While this shows that distance may not be a significant factor in transmission, it would be premature to draw solid conclusions without first evaluating the potential limits of our study design. The results indicate that the distances between animals differ greatly, with numerous interactions taking place at distances greater than 2 meters. Furthermore, the frequent contacts occurring at these longer distances are linked to elements that reduce the precision of models using distance-related characteristics. We also noted that the uneven distribution of interaction distances, very few close-range interactions, hindered our capacity to identify impacts on virus transmission. Future studies should focus on better assessing the effective distance for contamination, either by implementing numerical simulation for the trajectory of droplets, or by considering experimental infections at predetermined fixed distances.

Another interesting finding that stands out from the results reported earlier is the estimation of the minimum exposure period required for successful contamination. Our analysis reveals significant insights into the relationship between distance and contamination risk. The results indicate a positive correlation between exposure time and the inter-individual distance between susceptible-seed pairs. We compared the performance of the baseline model, which ignores distances even when individuals are moving, and the hypothetical case when animals are kept restrained at a fixed distance. In this case we used results from the Stratified 1 and Stratified 4 models, which explicitly include all the different ranges of distances. Results from these hypothetical cases models show that less exposure time (between 2/3 and 1/10 of the baseline case) is needed to reach a probability of transmission of 0.95 than the baseline model. This difference indicates that the integration of the proximity distance in the models improves the model estimates and helps be closer to real-world settings. Alternatively, considering that shorter distances between animals correspond to higher densities, we can infer how exposure times vary with density. The minimal exposure time scale of a factor 4 from a highly dense herd (distance < 0.5 m, density = 5 animals per m²) to a herd that is almost 35 less dense, or a factor 12 when we consider animals that can randomly move in a space 100 times larger. These findings strongly support the effectiveness of physical distancing measures and provide valuable insights into safety protocols in controlled environments. While future research should focus on reducing uncertainty ranges and investigating additional variables affecting exposure times, our current results offer a robust foundation for developing evidence-based safety guidelines and risk management strategies.

Direct animal contacts between virus excreting animals with susceptible animals are certainly the main way, or important way, for PPRV to spread but it might not be the only one. The virus might spread through shared water or feed or through feces, as well as urine as indicate by some authors [17,50]. But those possibilities have yet to be proved experimentally. Current studies have only detected genetic material and antigens in feces and urine, without direct evidence of infectious particles, despite frequent correlation between genetic material and viral particles. Studies of market dust samples have only documented genetic detection, highlighting the need for comprehensive viability assessment. The virus that spreads in shared spaces needs broad disease control that includes both the surroundings next to livestock use of common areas and items. Combining data captured with UWB together with camera recordings could provide more information about “hotspots” of transmission and the role of direct and indirect contact. A clear view of these facts shapes PPRV control plans and the implementation of biosecurity measures to stop transmission.

## Conclusion

Ultra-wideband (UWB) captors proved effective for monitoring goat movements and estimating infectious pathogen transmission probability and exposure time based on proximity data. The presence of viral transmission in communal environments necessitates broad control measures addressing both immediate livestock surroundings and shared resources. The current study presents certain limitations, including restricted sample size and controlled experimental conditions that may not fully reflect field scenarios. Environmental factors and seasonal variations were not extensively examined, potentially influencing transmission dynamics in ways not captured by our models. Future research might investigate the following: conditions marketplaces where animals stay in close contact with important movements, possibilities of indirect transmission routes through feed and water, duration of virus viability in excretions. Additional research should focus on optimizing proximity data collection methodologies and quantifying close contact frequencies to enhance modeling accuracy. Studies incorporating larger herd sizes, and field conditions across diverse geographical regions would validate the applicability of these findings. Additionally, investigation into environmental persistence of PPRV under varying climatic conditions would contribute valuable data for comprehensive control program development.

## Materials and methods

The study was conducted between June 2021 and June 2022. Five sessions of experiments were conducted during this period in June, August 2021, and January, April, and June 2022. In each session, two or three batches of six or seven animals were experimentally infected (exposure phase) and monitored. Four durations of the exposure phases were tested: 1 hour, 6 hours, 24 hours, and 44/48 hours. A total of twelve experiments were carried out for a total of 82 animals involved. The study was designed to detect 80% of the event of a transmission, if any, with a probability as low as 0.03. For the ease of the analysis, data from batches were classified using the session number (1 to 5), the batch number in each session (1 to 3), and the duration of the exposure phase (1h, 6h, 24h, 44h, 48h).

### Virus

In each experiment, a single animal (called the seeder) was infected with the PPRV Senegal 20-GP strain. The virus was isolated in BTS cells, a recombinant CV1 cell expressing the bovine SLAM protein (C. Adombi and A. Diallo, unpublished data), similar to the CHS cell, the CV1 cell expressing the goat SLAM [32], which is efficient for PPRV in vitro isolation. This virus was grown in the BTS cells, and the collected virus suspension was aliquoted and stored in a 1.5 ml tube at -80°C. It was used at the 3rd passage in the BTS cells. The contents of three tubes were defrosted and titrated independently by the endpoint dilution assay on the BTS cells in 96-well flat-bottom plates for their cytopathic effect (CPE). The virus suspension titer was expressed as the 50% tissue culture infectious dose (TCID50), calculated according to the Spearmann-Kärber methodology as described by [33]. The mean value of the three titrations was 10^3.2^ TCID50/ml. It belongs to the PPRV lineage IV according to its partial Np sequence data [40]. In preliminary animal pathogenicity testing (Diallo A., Diop. M, Sagna A. and M. Lo, unpublished data), all goats inoculated with this virus exhibited acute PPR symptoms: high fever up to 40-41°C, diarrhea, significant ocular and nasal discharges, and death in four out of ten inoculated animals.

### Animals

A total of 82 goats, naïve and aged between six and twelve months, were used in the twelve experiments. They were purchased from different farms in Senegal by a team from the Laboratoire National d’Elevage et de Recherches Vétérinaires (LNERV) of the Institut Sénégalais de Recherches Agricoles (ISRA). Before purchase, all were proved negative for both PPRV and antibodies anti-PPRV by the ID.vet Innovative Diagnostic PPR pen-side test (ID Rapid® PPR Antigen) and the ID.vet Innovative Diagnostic ID Screen**®** PPR Competition respectively. The tests were performed by the ISRA laboratory team in the village of the farmers before purchasing the animals. After the purchase, the animals were brought to the ISRA experimental farm at Sangalkam (14.7955°N, 17.2289°W; WGS 84), about 30 km from Dakar. Upon arrival they were housed in a 300 m² barn for a minimum of two weeks for quarantine and acclimatization. During the first and second weeks, they were submitted to a second round of antibody anti-PPRV detection assay (ID.vet Innovative Diagnostic ID Screen® PPR) and the Reverse Transcription Polymerase Chain Reaction (RT-PCR) for PPRV nucleic acid detection according to the method of [51]. Only animals that tested negative again in both assays were included in the experiments. Any goat that was tested positive by one of the two tests was immediately removed from the group and excluded from the study.

### Experimental facilities

At the Sangalkam experimental farm, the experimental infection protocol was conducted in a dedicated, specially equipped area of 40m² (Fig 8), located at about 50 meters from the quarantine barn. Within this area, paddock 2 was designated for controlled PPRV infection, while the animal-to-animal PPRV transmission experiments were conducted in paddock 1. Both open experimental barns were equipped with fenced pens designed to house each animal individually after the transmission experiments, to facilitate individual monitoring of PPR symptoms. The storage rooms were designated for storing animal feed and technical equipment, including syringes, blood collection tubes, gloves, coats, and containers for pathological sample collection. Outside the periods of experimental infection protocols, the animals were housed in the 300 m² barn. All barns, paddocks, and individual pens were equipped with troughs to provide animals with ad libitum access to feed and water. All the animal facilities were cleaned daily.

**Fig 8.**
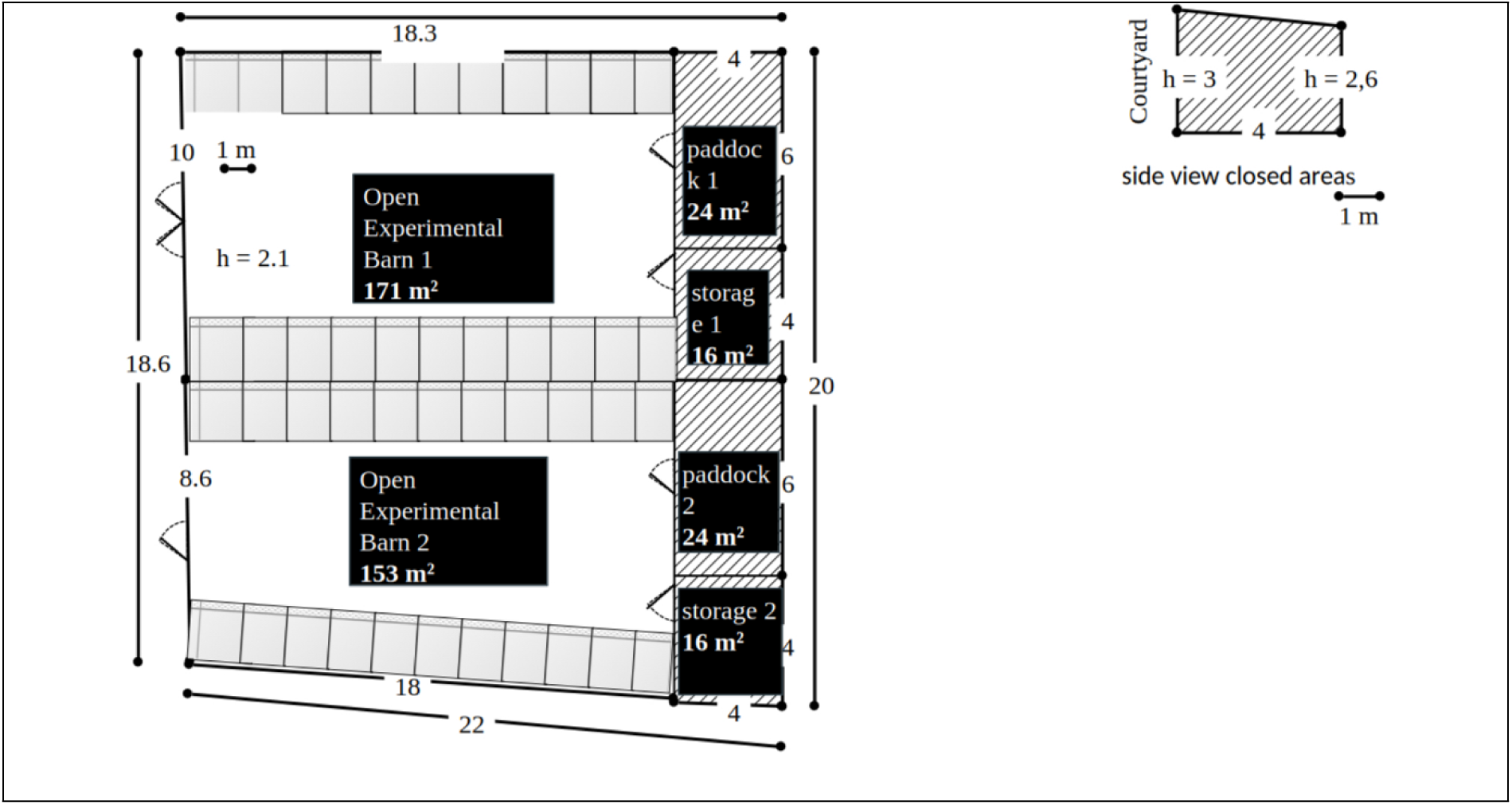
Schema of the Experimental Facility. This diagram shows the experimental facility maps including dimensions.

### Experimental design

Each experience was divided into three phases as shown in Fig 9:

1. Experimental Infection phase: infection of animal to be used for transmission of the PPRV to naïve animals by contact: the seeder.
2. Exposure Phase: The seeder is in contact with other naïve animals to allow transmission from animal-to-animal in a paddock for a period of variable duration.
3. Monitoring phase: exposed animals are isolated and monitored until they show symptoms or positive results to PPR.

**Fig 9.**
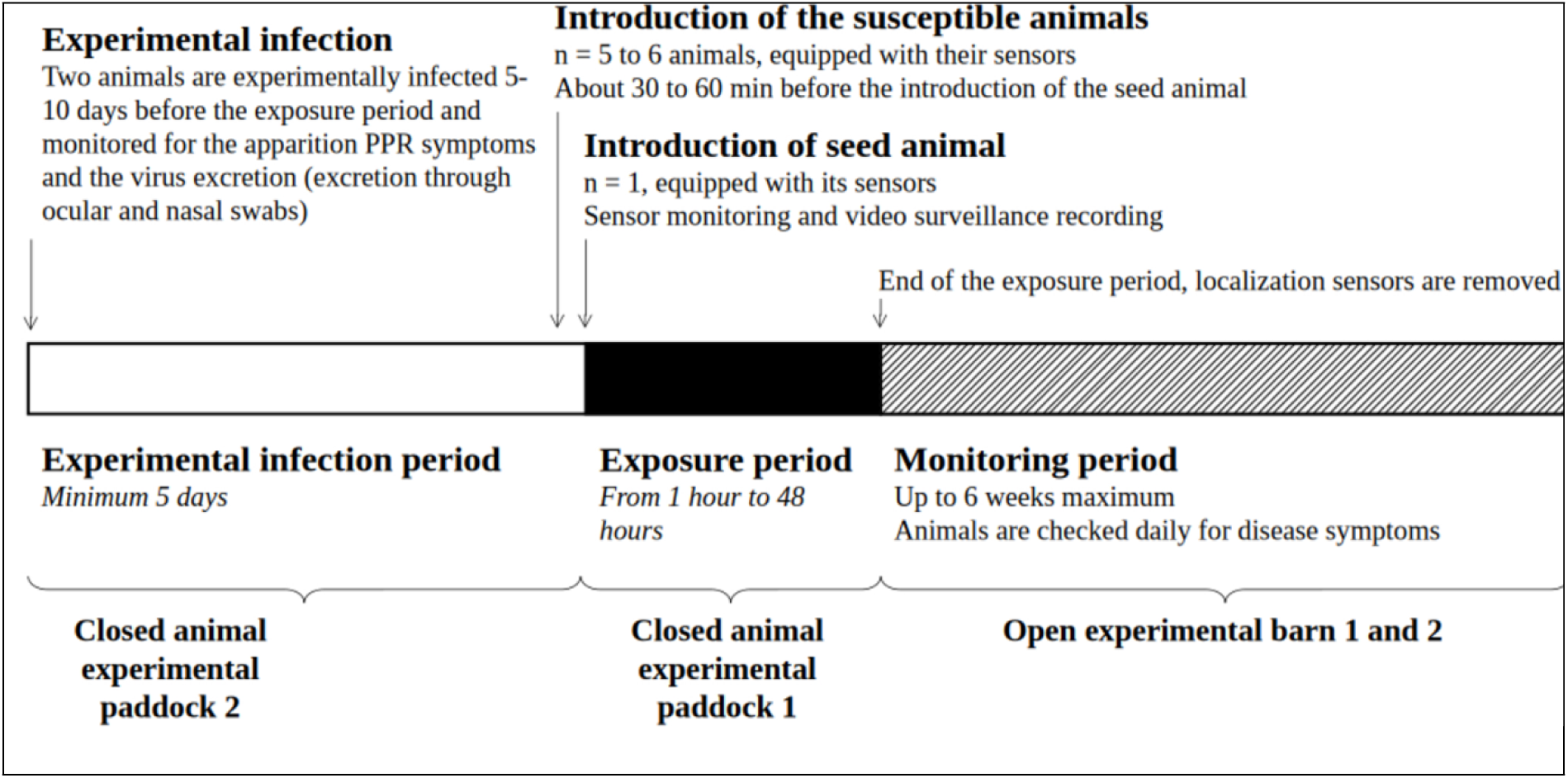
Detailed Schema of the Experimental Infection Protocol for Studying PPR lineage IV Transmission Dynamics. This diagram outlines the sequential stages of the experimental protocol designed to investigate the transmission of PPR among goats. The protocol includes three primary periods: the experimental infection period, the exposure period, and the monitoring period.

#### Experimental infection phase

In the preliminary part of the experiment, two goats were retrieved from the quarantine barn and moved to paddock 2 where from each animal ocular and nasal swabs were collected. Those collected samples will be used as “day 0” samples in different subsequent tests: antigen, nucleic acid, and serum detection tests. Just after this procedure, the two animals were subcutaneously inoculated with 10^3^ TCID50 of PPRV Senegal 20-GP in 2 ml. The subjects were then housed in paddock 2 and subjected to daily clinical observation for monitoring any clinical signs of PPR expression, including rectal temperatures. All observations were recorded on a dedicated clinical sheet. The ocular and nasal swabs were also collected daily and submitted to tests for PPRV antigen detection and the nucleic acid detection by real-time RT-PCR. Following preliminary results with that test, a good level of virus excretion in the swab is noted when the threshold cycle number (CT value) is less than 26. So once an inoculated animal expressed clinical signs of PPR and a good level of PPRV excretion in the collected swabs based on their test results, it was removed from paddock 2 to paddock 1 to serve as a seeder for the exposure phase. Usually, this period of experimental infections lasted between six to ten days post infection.

#### Exposure phase

Once the seed animal was identified, five/six animals from the quarantine barn were randomly selected for the exposure period and housed in paddock 1. Each of the selected animals was bled for the day 0 serum collection. This bleeding was carried out along with the ocular and nasal swabs (in duplicate) collection. All those samples will serve as day 0 samples in the tests of other subsequently collected samples testing. Each animal was equipped with a collar-mounted UWB sensor (BeSpoon - STMicroelectronics, France) to triangulate their relative positions and estimate distances all along the experiment. At the time of the introduction of the seed animal into paddock 1, recordings were initiated. The length of exposure of the naïve animals varied, with each exposure duration representing a distinct experiment.

#### Monitoring phase

At the end of the exposure phase, each animal was retrieved, its UWB collar was removed, and it was placed in an individual fenced pen in either Open Experimental Barn 1 or 2. This marked the start of the monitoring period that lasted for about 5 weeks (except for experiment 4 – 48 hour exposure – for which the duration was 8 weeks). The health status of each animal was monitored daily through visual observations without touching them. Once a week the animals were handled for bleeding for serum and nasal/ocular swabs collections. It was only on that occasion the animals were touched and only by one animal attendant and the scientist in charge of the trial. For that, they wore personal protective equipment (PPE) which were changed between each animal. Outside of these sampling periods, only the animal attendant was allowed to enter the pens for cleaning and for providing feed and water in dedicated troughs, without making direct contact with the animals.

At any time during the monitoring phase, once an animal looked sick, its rectal temperature was recorded, nasal and ocular swabs were collected along with the blood for serum. In such cases, only apparent sick animals were handled. In the event of death, a post-mortem autopsy of the animal was performed with the collection of some pathological samples: lymph nodes and piece of lung to be tested for the PPRV nucleic acid detection by real-time RT-PCR. The findings were recorded.

### Data collection for indoor positioning

All animals in the experiments were equipped with UWB devices to monitor and record, in near-real time, the precise indoor location of everyone. UWB solutions are used to measure the distance between a reference sensor (referred to as an anchor) and a sensor whose position is to be determined (referred to as a tag, equipped on the animal) using different metrics. The theoretical performance of the solution allows positioning of an object with an accuracy of +/- 20 centimeters, but these performances depend notably on the tracking environment and the deployment conditions. For these experiments, paddock 1 was equipped with six anchors (UWB BeSpoon) fixed to the ceiling, one at each corner and two near the center of the room. To ensure accurate indoor positioning, the 3D position of each anchor was mapped relative to one corner of the room. Each anchor was powered by a Power Over Ethernet cable connected to a dedicated 16-port switch (DES-1018MP; D-Link, Taipei - Taïwan). The tags (BeSpoon Industrial Tag; Length: 101mm, Width: 52mm, Thickness: 25mm; BeSpoon - STMicroelectronics, France) were enclosed in a custom 3D-printed case to provide shock protection and ensure better attachment to the collar. The UWB system was controlled via a dedicated local server (a laptop with a Linux operating system), which provided a user interface for verifying the general system status and initiating data acquisitions. The visualization interface allowed for the definition of tracking zones and the configuration of data recording. For our experimental needs, the acquisition frequency was set to f = 1 Hz (one acquisition per second). Data were stored in a dedicated folder on the server at the end of each tracking session or, by default, every 24 hours. The data is stored in a.csv file, with each line representing the estimated position of each tag on the x, y, and z axes. Distances between the infected individual and each susceptible animal were then computed from these data, using only the x and y axes. This frequency of indoor positioning provided a comprehensive temporal map of animal interactions. High-resolution proximity data enabled us to analyze not only the frequency of close contacts, but also the duration of these interactions at different distance thresholds.

### Network analysis of the contact pattern

Data collected from UWB devices were used to reconstruct the contact patterns between animals. A temporal network approach was applied to represent and analyze these interactions. In this framework, animals are represented as nodes, and their interactions (based on proximity) as links with varying strengths (in this case, the distance). In a temporal network, the number of nodes, links, and their strengths (i.e., distance) can change over time.

In each experiment, the network consisted of 6 or 7 nodes—corresponding to the number of animals in the pen—and 15 or 21 possible links, respectively. While the number of nodes and links remained constant each time, the distances between animals varied. For each experiment, we analyzed the distribution of these distances. The Kruskal-Wallis test, followed by pairwise Wilcoxon tests, was used to assess whether significant differences existed among the distance distributions.

Additionally, we used the method developed by Berlingerio et al. [52] to identify possible patterns among animals. This approach relies on the use of a particular form of the Jaccard measure for weighted networks to quantify similarity over time:

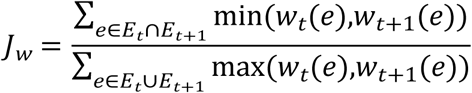

where *e* indicates a link on the network *w_t_(e)* the corresponding weight, *E_t,t+1_* the set of edges at time *t* and *t+1*. After computing the Jaccard similarity between all pairs of consecutive time points, a hierarchical cluster approach is used to identify possible *eras,* period of times during which the dynamics among animals are different.

### Statistical analysis and models

We employed a Bayesian statistical framework to estimate the transmission rate, incubation period, and model transmission probability. Depending on the particular quantity to estimate, we used different likelihoods to characterize the binary outcome of infection status and the duration of the incubation phase. In all cases we used minimally informative prior distributions to minimize potential bias and applied the Markov Chain Monte Carlo (MCMC) approach to estimate model parameters.

#### Transmission rate

We estimated the transmission rate of the strain following the procedure of Dekker et al. [53]. The infection outcome for each susceptible individual in an interval of time Δt modeled as an independent and identically distributed Bernoulli random variable with infection probability *p_t_*.

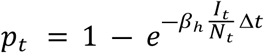

Where *N_t_* is the total number of animals in the pen/paddock, *I_t_* the number of initially infected individuals (1 in all cases) and Δt is the duration of the exposure phase ranges from 1 hour to 48 hours (Δt = 1, 6, 24, 44, 48). Due to the variation in the length of the exposure phase, we estimated *β_h_* the transmission rate per hour. We estimated *β_h_* using the No-U-Turn Sampler (NUTS), fitting the model to serological data using a binomial likelihood.

#### Basic reproductive number

In a continuous transmission model, considering an exponential process, the daily transmission rate *β* can be estimated as:

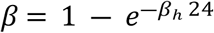

Based on this, a first estimate of the basic reproductive number R_0_ was calculated using the following formula derived from the SIR model definition:

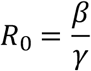

where, β is the daily transmission rate, and γ stands for the recovery rate commonly used in modeling works for PPR.

#### Incubation period estimation

Following the same procedure that was adopted by Miura et al. [54], we estimated the incubation period by fitting a parametric distribution to the animal follow-up data. In order to estimate the incubation period, one of the following information were taken into consideration for each animal when available: the day on which symptoms were first observed, the day on which a positive result was obtained from the PCR test (indicating the presence of the virus), and the day on which a positive result was obtained from the serology test.

We used data on the time at which symptoms first appeared, or on positive RT-PCR or serology results, to estimate the incubation period. After a preliminary analysis to check if exposure duration could impact the length of the incubation period survival analysis was conducted to estimate the average incubation period disregarding categorization in exposure groups. To take account of the uncertainty of the moment of infection occurrence during the moment of exposure phase, this was randomly extracted for each animal. Three typical parametric distributions used in survival analysis (Weibull, lognormal, and Gamma) were used for likelihood and chose the one with the lowest widely applicable information criterion (WAIC).

#### Transmission probability

Our analytical approach comprised four distinct modeling strategies for PPRV transmission in our study: (i) baseline model, (ii) segmented model, (iii) envelope model, and (iv) Stratified model. In our Bayesian statistical approach, we initially developed a baseline model that intentionally excluded proximity factors, focusing solely on temporal exposure patterns (see more details in Fig 10). This approach allowed us to first characterize transmission dynamics without spatial considerations. Subsequently, we progressively incorporated proximity data from UWB to systematically evaluate whether distance information significantly enhances transmission risk prediction. By comparing models with increasing spatial complexity, from a null temporal model to models integrating precise inter-animal distances, we aimed to empirically determine the statistical significance of proximity in PPRV transmission mechanisms. In all these models, we used Bernoulli likelihood to consider individual differences in contact patterns and activity.

**Fig 10.**
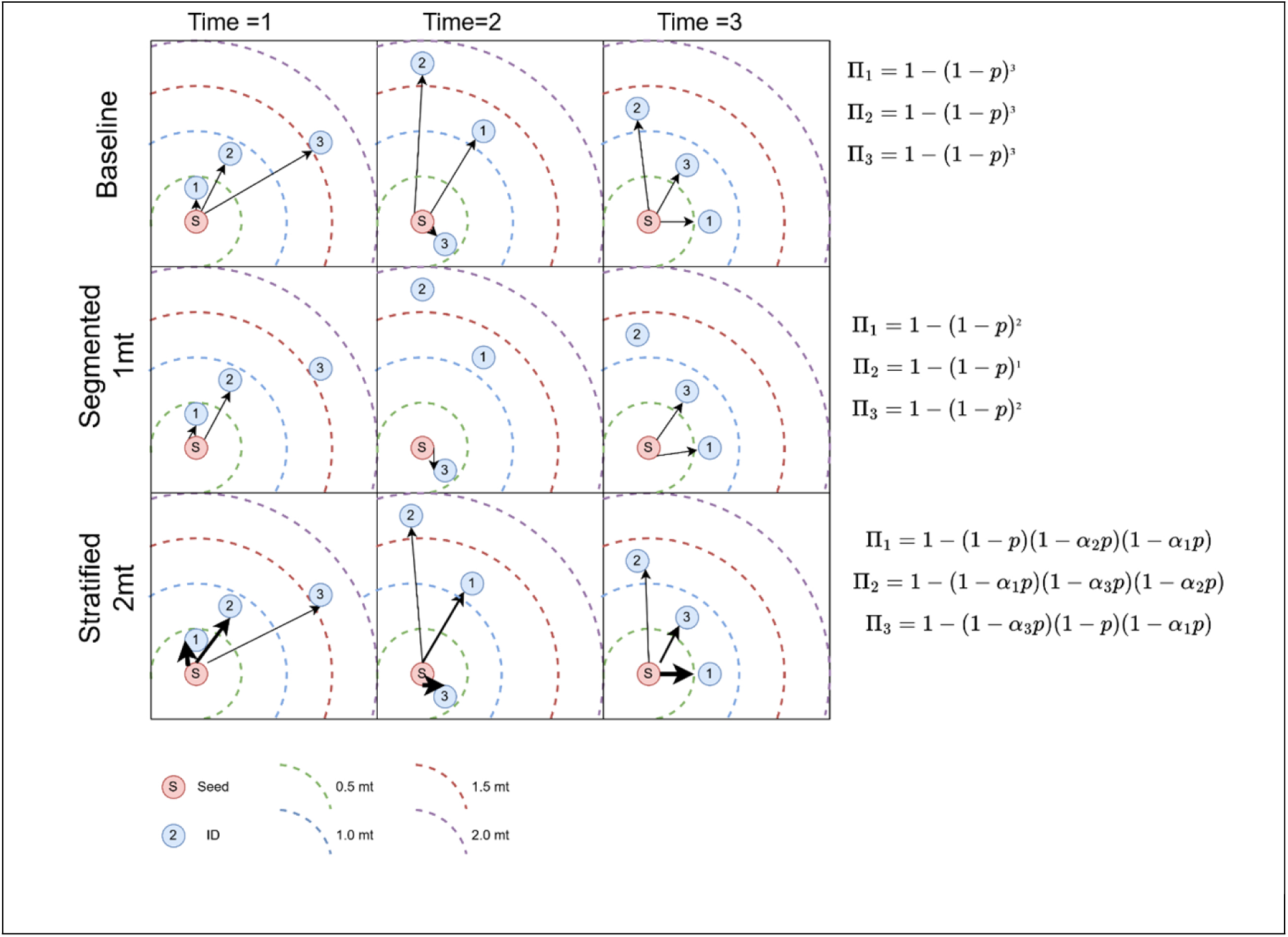
Graphical representation of three different models (Baseline, Segmented, and Stratified) illustrating the spread of infection over time. The infected seeder individual, denoted as S, interacts with susceptible individuals (1), (2), and (3). The arcs represent the varying distances between S and the susceptible individuals at three time points: t = 1, t = 2, and t = 3 (in minutes). The circles depict the dynamic radius of interaction, which either increases or decreases over time, showing the changing proximity and potential for infection spread in all orientations. The envelope model is a modification of Stratified model, where ranges are determined based on animals’ movement patterns.

##### Baseline model

Our baseline model, which serves as the null hypothesis, assumes a constant probability of transmission per unit of time, independent of proximity. The individual cumulative probability of infection *P* for this model is expressed as follows:

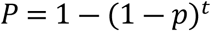

where *p* represents the probability of a successful transmission per unit time, and *t* the total duration of the experiment. The simplicity of this model provided a fundamental benchmark against which more complex models could be compared.

##### Segmented model

Building on the baseline approach, our segmented model has incorporated distance-based stratification to account for the potential influence of proximity on the virus transmission probability. This model uses the same fundamental equation as the baseline model:

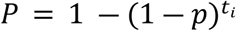

However, in this context, *t_i_* represents the cumulative time spent within specific distance thresholds (1 m, 1.5 m, and 2 m). Separate models were fitted for each threshold, enabling us to discern how the probability of transmission might vary with different distances.

#### Stratified model

Our stratified model represents a more subtle method of integrating transmission dynamics as a function of distance. The observed distances were divided into different ranges:

- *t_1_*: cumulative time spent between 0.05 m and 0.5 m,
- *t_2_*: cumulative time spent between 0.5 m and 1 m,
- *t_3_*: cumulative time spent between 1 m and 2 m,
- *t_4_*: cumulative time spent beyond 2 m.

The probability of infection for this model is given by:

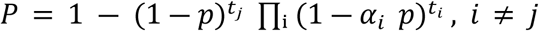

where *p* is the probability of success, *α_i_* are scale factors for different distances, and *t_i_*, *t_j_*represents the time spent in each range. This formulation allows for a differential weighting of transmission risk according to proximity. We studied five variants of this model, each emphasizing different distances:

- Stratified 1: *j* = 1, *i* = 2, 3, 4
- Stratified 2: *j* = 1, *i* = 2
- Stratified 3: *j* = 3, *i* = 4
- Stratified 4: *j* = 2, *i* = 3, 4
- Stratified 5: *j* = 1, *i* = 2,3

##### Envelope model

Our most sophisticated approach, the envelope model, integrates both temporal and spatial proximity data clustering. We identified distinct envelope interaction groups for each individual and used the total duration of these groups along with their median distances. In this model, the probability of an individual’s infection is expressed as follows:

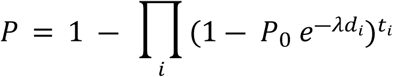

where *P_0_* represents the maximum probability of success when the distance is equal to zero, λ a decay rate, *d_i_* is the median distance of envelope *i*, and *t_i_* is the cumulative time in envelope *i*. This model was applied with different distance filters (≤ 1 m, ≤ 1.5 m, and ≤ 2 m), with distinct models adjusted for each filter.

### Parameter estimation

The choice of prior distributions for model parameters was guided by both biological plausibility and computational considerations. The specific parametrizations of the priors were selected based on preliminary analyses and expert knowledge of PPRV transmission dynamics. Since individuals could vary based on their activity patterns, we used a binomial likelihood to elicit the differences among individuals’ patterns.

We used the NUTS, an MCMC method. This approach facilitated efficient exploration of the parameter space and estimation of posterior distributions. Our sampling protocol was designed to ensure thorough exploration of the parameter space. We configured the sampler to draw 10,000 samples per chain, with a tuning phase of 1,000 iterations, and ran ten independent chains. This configuration allowed for comprehensive sampling and robust convergence diagnostics. We chose a high target acceptance rate of 0.95 to promote efficient exploration of parameter space, which was particularly important for our more complex models. Posterior distributions were obtained for each model parameter, and the Bayesian estimates were summarized using the posterior mean estimate and the 95% highest (posterior) density interval (HDI). The 95% HDI is the minimum Bayesian credible interval (BCI) that contains 95% of the posterior probability distribution. Convergence of the chains was confirmed by visual inspection of trace and density plots and, the Gelman—Rubin diagnostic (R^) which compares the within-chain and between-chain variances [55]. If the chains have converged, the variances should be similar and results in an R^ close to 1.

### Model evaluation

Model comparison was facilitated through the computation of WAIC [56] because it provides an estimate of predictive accuracy while accounting for model complexity, without requiring cross-validation or separate training/test datasets. WAIC estimates how well a model generalizes to unseen data by balancing goodness of fit (how well the model fits the observed data) and penalizing complexity (to avoid overfitting). A model with a lower WAIC value is preferable, as it indicates better predictive performance. However, models with close WAIC values can be considered as having similar performance, and other factors such as interpretability and model simplicity can be considered for the final selection.

We conducted a receiver operating characteristic (ROC) curve analysis and generated areas under the curve (AUCs) estimates using the individual transmission probability for each model to compare their binary classification performance. The ROC curve is a graphical representation that plots the True Positive Rate (TPR) against the False Positive Rate (FPR) across all possible thresholds. The AUC values range from 0 to 1, with the higher AUC values signifying superior performance of the predictive model.

We assessed the goodness of fit through posterior predictive checks using aggregated Bernoulli draws at the experiment level. This approach involves simulating replicated datasets by drawing from a Bernoulli distribution and then summing up these binary classifications to compute the total number of positive outcomes. We compared the mode of the simulated aggregates to the observed number of positive cases for each experiment graphically.

The Bayesian analyses were performed using the MCMC algorithm and were conducted in Python (version 3.9) using the PyMC language model (version 5.10.3) with BlackJax backend (version 1.1.0). WAIC computations were handled using the built-in function from Arviz library (version 0.17.0). The statistical tests were conducted using R (version 4.4.2).

## Acknowledgments

The authors are grateful to Mr. Abdel Sakaly Sagna and Mr. Djiby Ka for their technical support during the implementation of the animal experiments.

## Ethics statement

All the animal experiments have been carried out by staff of the LNERV/ISRA at the ISRA experimental farm. LNERV/ISRA is the Senegalese national veterinary laboratory for diagnostics and research in animal health and production. The procedures and protocols of the animal experiments which results have been reported here were submitted to and approved by/ the Scientific Direction of LNERV/ISRA, a competent body for animal research.

## Code and data availability

The code and data used in this study are publicly available on GitHub at https://github.com/mnhili/PPR_lineageIV_transmission.

## Supporting information

**S1 File. Supplementary information**. This document file contains additional analyses and extended methodological details supporting the findings of the study.

**S2 File. Official dataset**. This Excel file contains follow-up data from experimentally infected goats, organized by individual experiment.

